# Bayesian Model Selection Maps for group studies using M/EEG data

**DOI:** 10.1101/365056

**Authors:** Clare D. Harris, Elise G. Rowe, Roshini Randeniya, Marta I. Garrido

## Abstract

Predictive coding postulates that we make (top-down) predictions about the world and that we continuously compare incoming (bottom-up) sensory information with these predictions, in order to update our models and perception so as to better reflect reality. That is, our so-called ‘Bayesian brains’ continuously create and update generative models of the world, inferring (hidden) causes from (sensory) consequences. Neuroimaging datasets enable the detailed investigation of such modelling and updating processes, and these datasets can themselves be analysed with Bayesian approaches. These offer methodological advantages over classical statistics. Specifically, any number of models can be compared, the models need not be nested, and the ‘null model’ can be accepted (rather than only failing to be rejected as in frequentist inference). This methodological paper explains how to construct posterior probability maps (PPMs) for Bayesian Model Selection (BMS) at the group level using electroencephalography (EEG) or magnetoencephalography (MEG) data. The method has only recently been used for EEG data, after originally being developed and applied in the context of functional magnetic resonance imaging (fMRI) analysis. Here, we describe how this method can be adapted for EEG using the Statistical Parametric Mapping (SPM) software package for MATLAB. The method enables the comparison of an arbitrary number of hypotheses (or explanations for observed responses), at each and every voxel in the brain (source level) and/or in the scalp-time volume (scalp level), both within participants and at the group level. The method is illustrated here using mismatch negativity (MMN) data from a group of participants performing an audio-spatial oddball attention task. All data and code are provided in keeping with the Open Science movement. In so doing, we hope to enable others in the field of M/EEG to implement our methods so as to address their own questions of interest.

## 1 Introduction

The statistical testing of hypotheses originated with Thomas Bayes (Neyman and Pearson, 1933), whose famous eponymous theorem (Bayes and Price, 1763) can be written in terms of probability densities as follows:

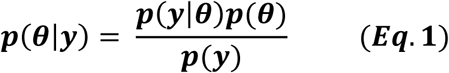

where *θ* denotes unobserved parameters, *y* denotes observed quantities, and *p*(θ*|*y) denotes the probability *p* of the unknown parameters θ, given (“|”) the set of observed quantities *y*. More generally, *p*(event|knowledge) denotes the probability of an event given existing knowledge. In other words, Bayes conceptualises statistics as simply the plausibility of a hypothesis given the knowledge available (Meinert, 2012).

Bayes’ theorem allows one to update one’s knowledge of the previously-estimated (or “prior”) probability of causes, to a new estimate, the “posterior” probability of possible causes. This process can be repeated indefinitely, with the prior being recursively updated to the new posterior each time. This gives rise to multiple intuitive and useful data analysis methods, one of which is the explained in detail in this paper.

Even when it first appeared, Bayes’ theorem was recognised as an expression of “common sense,” a “foundation for all reasonings concerning past facts,” (Bayes and Price, 1763). Centuries later, neuroscientific evidence suggests that Bayes theorem may not only explain our “common sense” and internal reasoning processes, but may be common to all our senses: it can actually explain the way in which we use our various senses to perceive the world. That is, Bayesian statistics can be used to accurately model and predict the ways in which our own brains process information (Dayan et al., 1995; Feldman and Friston 2010; Friston, 2012; Hohwy, 2013). This has given rise to the concepts of predictive coding and the Bayesian brain. In this context, it is unsurprising that Bayesian approaches to statistics have high face validity (Friston and Penny, 2003). This allows for intuitive descriptions of probability and enables experimental results to be relatively easily understood and communicated both within and between scientific communities, as well as to the general public (Dunson, 2001).

Despite the intuitiveness of Bayesian approaches, however, the mainstay of hypothesis-testing since the twentieth century (Vallverdú, 2008) has instead been classical or frequentist statistics, which conceptualises probability as a ‘long-run frequency’ of events, and which has dominated most approaches to neuroimaging analysis to date (Penny et al., 2007). For example, creating statistical parametric maps (SPMs), which is a popular method of analysing neuroimaging data, mainly involves frequentist approaches (Friston and Penny, 2003).

In frequentist statistics, the null hypothesis (that there is no relationship between the causes and the data) is compared with one alternative hypothesis; the null is then either rejected in favour of the alternative hypothesis, or it fails to be rejected – it can never be directly “supported.” Rejection of the null depends on the somewhat unintuitive p-value, which communicates how likely it is that the effect (of at least the size seen in the experiment), would be seen in the absence of a true effect, if the experiment were repeated many times. This is a more complex and counterintuitive way of communicating results compared to Bayesian statistics (where the probability of the hypothesis in question is what is being estimated and communicated). Also, unfortunately, multiple different models cannot be compared at once, and the null and the alternative models need to be nested for frequentist statistical tests to be feasible (Rosa et al., 2010). These features cause frequentist statistics to be less useful in certain contexts, compared to the approaches enabled by Bayesian statistics.

In recent decades, Bayesian approaches are becoming increasingly recognised for their superior utility for addressing certain questions and in specific data analysis situations, as explained below (Beal, 2003; Rosa, et al., 2010; Penny and Ridgway, 2013). Importantly, with Bayesian approaches to data analysis, any number of models can be compared, the models need not be nested, and the ‘null model’ can be accepted (Rosa et al., 2010). The fact that Bayesian hypothesis-testing also allows researchers to evaluate the likelihood of the null hypothesis is crucially important in light of the replication crisis in psychology and neuroscience (Hartshorne, 2012; Larson and Carbine, 2017; Szucs et al., 2017). Importantly, results supporting the null hypothesis are equally noteworthy or reportable as other results within Bayesian statistics. The use of Bayesian statistics may also ameliorate some statistical power-related problems documented in the literature (Dienes, 2016).

Even though Bayesian statistics has gained popularity in the context of ‘accepting the null’, its strength lies beyond this, in the sense that it enables the relative quantification of any number of *alternative* models (or hypotheses). In Bayesian Model Selection (BMS), models are compared based on the probability of observing a particular dataset given each model’s parameters. The probability of obtaining observed data, y, given model *m, p(y|m)*, is known as the model evidence. In BMS, an approximation of the model evidence is calculated for multiple models; the model evidences are then compared to determine which model returns the highest probability of generating the particular dataset in question (Rosa et al., 2010).

A computationally efficient and relatively accurate (Penny et al., 2009) method of approximating the model evidence is to use variational Bayes (VB). If each participant in the dataset is assumed to have the same model explaining their data, then this is called a fixed effects (FFX) approach. If, on the other hand, every participant is permitted to have their own (potentially different) model, this is called a random effects (RFX) approach.

An elegant approach to succinctly communicating results is to use Posterior Probability maps (PPMs), which provide a visual depiction of the spatial and/or temporal locations in which a particular model is more probable than the alternatives considered, given the experimental data in question. The development of PPMs is essentially the Bayesian alternative to the creation of SPMs (Friston and Penny, 2003). PPMs may display the posterior probability of the models (the probability that a model explains the data), or, alternatively, they may be displayed as Exceedance Probability Maps (EPMs), which are maps of the probabilities that a model (say *k*) is *more* likely compared to all other (*K*) models considered (Rosa et al., 2010). (EPMs will be identical to posterior probability maps in cases where there are only two models being considered, as in this study.) EPMs are useful in that they allow us to directly quantify which model is more probable than the other/s considered.

The data analysis method that forms the focus of this paper is Posterior Probability Mapping with an RFX approach to VB. First introduced (Rosa et al., 2010) for functional magnetic resonance imaging (fMRI), the method has recently been adapted for inference using electroencephalography (EEG) data (Garrido et al., 2017). In their study, Garrido and colleagues (2017) used variational Bayes to approximate the log of the model evidence for each voxel (in space and time) in every participant, in order to construct PPMs at the group level. They did this in the context of comparing between two computational models describing the relationship between attention and prediction in auditory processing. While that paper focused on using this Bayesian methodology to address an important neuroscientific question, the precise way in which Rosa and colleagues’ (2010) methods were adapted for use with EEG data have not been formally described to date – leading to the purpose of this paper.

Here, we describe in a tutorial-like manner how to build and compare PPMs for EEG and/or magnetoencephalography (MEG) data (M/EEG), using an RFX approach to VB. This approach provides useful ways of displaying the probabilities of different models at different times and brain locations, given any set of neuroimaging data (as done in (Garrido et al., 2017)) using the Statistical Parametric Mapping (SPM) software package for MATLAB. Furthermore, in keeping with the Open Science movement, we provide the full EEG dataset (https://figshare.com/s/1ef6dd4bbdd4059e3891) and the code (https://github.com/ClareDiane/BMS4EEG) to facilitate future use of the method. In so doing, we hope that this paper and its associated scripts will enable others in the field of M/EEG to implement our methods to address their own questions of interest.

## 2 Theory

In frequentist hypothesis testing, what is actually being tested is the null hypothesis (i.e. that there is no relationship between the variables of interest; Friston, 2007b). If it is assumed that there is a linear relationship between the causes and data, then the relationship between the causes (*x*) and data (*y*) can be represented as below (Friston, 2007b):

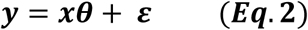

where *y* denotes data, *x* denotes causes and *ε* is an error term. The null hypothesis is that the relationship between the causes and data does not exist, that is, *θ* = 0. The null hypothesis is compared to one alternative hypothesis; the null is then either rejected in favour of the alternative hypothesis, or it fails to be rejected – it can never be directly “supported.”

Using the frequentist framework, one cannot test multiple models at once (unlike what can be done when using Bayesian approaches). (In this setting, a model corresponds to a particular mixture of explanatory variables in the design matrix *x*.) Even if one only wishes to test one model against the null, however, frequentist statistics still gives rise to problems unless the null and alternate models are nested. When the variables in one model cannot be expressed as a linear combination of the variables in another model, the two models are said to be non-nested (McAleer, 1995). Non-nested models usually arise when model specifications are subject to differences in their auxiliary assumptions or in their theoretical approaches, and can still be dealt with by making specific modifications to frequentist approaches (McAleer, 1995; Horn, 1987). However, there are many situations where Bayesian approaches are more appropriate for non-nested models than adapted frequentist inference (Rosa et al., 2010). Indeed, Penny et al. (2007), showed that functional magnetic resonance imaging (fMRI) haemodynamic basis sets are best compared using Bayesian approaches to non-nested models (Penny et al., 2007).

Furthermore, Bayesian approaches to statistics have long been recognised for their relative advantages outside of the realm of neuroimaging. In clinical trials, Bayesian experimental design techniques and interim analyses have been found to improve trials’ statistical power, cost-effectiveness and clinical outcomes (e.g. Trippa et al., 2012; Connor et al., 2013), compared to when classical approaches are used alone. Bayesian statistics are also especially useful in the worlds of computational physics (Mohammad-Djafari, 2002) and biology (Needham et al., 2007), and in machine learning (Lappalainen and Miskin, 2000).

The aim of BMS is to adjudicate between models using each one’s *model evidence*. Also written as *p(y*|*m*), the model evidence is defined as the probability (*p*) of obtaining observed data (denoted *y*) given the model (denoted *m*). It is given by the following integral:

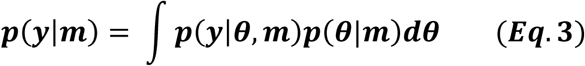

This integral is usually intractable, so numerous methods have been developed to approximate it. As Blei et al., (2017) succinctly summarise, there are two main ways to solve the problem of approximating the integral above. One is to sample a Markov chain (Blei et al., 2017), and the other is to use optimisation. The conversion of an integration problem into an optimisation problem is due to Richard Feynman, who introduced variational free energy in the setting of path integral problems in quantum electrodynamics (Feynman et al., 2010; Feynman and Brown, 1942). By inducing a bound on the integral above – through an approximate posterior density (please see below) – one converts an intractable integration problem into a relatively straightforward optimisation problem, that can be solved using gradient descent.

Some of the specific approximation methods that have been used to date include Annealed Importance Sampling (AIS; Neal, 1998; Penny and Sengupta, 2016), Bayesian Information Criterion (BIC) measures (Rissanen, 1978; Penny, 2012), Akaike Information Criterion (AIC) measures (Akaike, 1980; Penny, 2012), and finally, the variational Free Energy (F), which was first applied to the analysis of functional neuroimaging time series by Penny, Kiebel and Friston (2003) and which is explained in this paper (Rosa et al., 2010). These methods have varying degrees of accuracy and computational complexity, and have been studied in detail elsewhere (Beal and Ghahramani, 2003; Penny et al., 2004; Penny, 2012). The variational Free Energy provides a relatively high level of accuracy, without a great computational cost (Rosa et al., 2010), and so it is unsurprising that it is widely used in neuroimaging (Rosa et al., 2010). The Free Energy formula is (Penny et al., 2003):

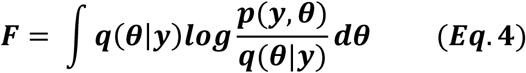

where *q(θ*|*y*) is an (initially) arbitrary distribution of the parameters *θ* given the data at each voxel *y, p(y*,*θ*) denotes the joint probability of the data and the parameters occurring, and d*θ* simply denotes that the integral given by F is with respect to the model parameters *θ*.

The “variational” term in variational Free Energy, and in variational Bayes (VB), refers to the branch of calculus (the calculus of variations) that deals with maximising or minimising functionals, or integrals. The utility of variational calculus in neuroimaging analysis has been reviewed in numerous other papers (Friston et al., 2008). In brief, the aim in variational Bayes is to maximise the functional given by the equation above. The reason for doing this is that it provides information about the model evidence. More specifically, the Free Energy relates to the log of the model evidence (or log-model evidence) as described by the following equation, known as the fundamental equation of variational Bayes (Penny et al., 2003):

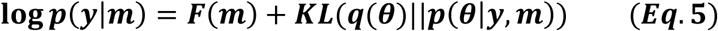

where log *p(y*|*m*) is the log-model evidence, *F* is the variational Free Energy, and 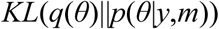 is the Kullback-Leibler divergence, or relative information, with respect to the approximate distribution *q(θ*) and the distribution that is diverging from it, namely the true distribution, *p(θ|y,m)*, as further described below.

The reason why Free Energy can be used as an approximation of the model evidence is better understood in light of the meaning of the second term in the fundamental VB equation, the Kullback-Leibler (KL) divergence (Penny et al., 2003). The equation for this is:

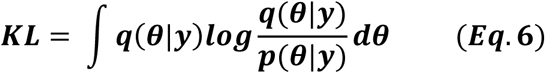

where all terms listed here have the same meanings as defined in earlier paragraphs. The KL divergence is also known as KL information, and this is because it is a measure of the information “difference” or divergence between two distributions. It can be derived by considering the so-called cross-entropy and entropy of the two distributions respectively, as outlined below (Carter, 2011). The concept of “relative entropy” is essentially “average information,” with “information” being defined as Shannon (1984) originally introduced:

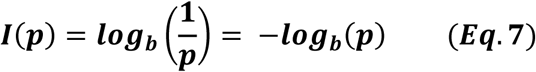

where *I(p)* is the information given by observation of an event of probability *p*, and 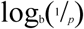 is the logarithm (in base *b*) of the inverse of the probability of that event. The formula above is used to derive the “average information,” also sometimes referred to a relative entropy, from a set of events. A related concept is the “cross entropy” between two distributions (see Carter, 2011); and the difference between the cross entropy and the entropy of the original/true distribution is equivalent to the KL divergence. Being a measure of information, the KL divergence has the property that it is non-negative; consequently, the lowest value it can take is zero.

The KL divergence between two distributions is zero only if the two distributions are equivalent. The closer KL is to zero, the less dissimilar the two distributions are. Thus, minimising KL is equivalent to maximising F, and F is said to provide a lower bound on the log-evidence. The aim of VB learning is to maximise F so that the approximate posterior thereby becomes as close as possible to the true posterior (Penny et al., 2007).

If (and only if) the KL divergence is zero, then F is equal to the log-model evidence. The free energy thus provides a *lower bound* on the log-evidence of the model, which is why iteratively optimising it allows us to proceed with BMS using F as an *approximation* of the log-model evidence (Penny et al., 2007). As the KL divergence is minimised by an iterative process of optimisation, F becomes an increasingly “tighter” lower bound on the desired (actual) log-model evidence; owing to this, BMS can proceed using F as a “surrogate” for the log-model evidence (Rosa et al., 2010). The iterations continue until improvements in F are very small (below some desired threshold). This method of estimating the log-model evidence is implemented in the second script described in the Implementation section (“BMS2_ModelSpec_VB.m”).

Although it has been summarised here, it is also worth noting that VB is further fleshed out in multiple other research papers (Penny et al., 2003; Friston et al., 2007; Friston and Penny, 2007; Penny et al., 2007) and tutorials (Lappalainen and Miskin, 2000). In *Statistical Parametric Mapping*, Friston (2007a) provides the mathematical derivations for the fundamental equation of variational Bayes, and his colleagues provide a full explanation of its application to BMS (Penny et al., 2007).

The application of VB in the context of fMRI analysis has been described in detail elsewhere (Rosa et al., 2010; Stephan et al., 2009; Penny et al., 2007). Penny and colleagues (2007) used Bayesian spatiotemporal models of within-subject log-model evidence maps for fMRI data, in order to make voxel-wise comparison of these maps and thereby to make inferences about regionally specific effects. Rosa and colleagues (2010) developed their approach by combining the methods described by Penny et al. (2007) with those of Stephan et al. (2009), who used an RFX approach to VB, as described below.

After the log-model evidence has been estimated as described above, given uniform priors over models, one can then estimate posterior model probabilities by comparing model-evidences between models. The ratio between model evidences, or Bayes factor (BF), can be used to estimate posterior model probabilities. A BF greater than 20 is equivalent to a posterior model probability greater than 0.95 (Kass and Raftery, 1995), which is reminiscent of the typical p-value smaller than 0.05. The product of Bayes factors over all subjects is called the Group Bayes Factor (GBF), and it gives the relative probability that one model (relative to another) applies to the entire group of subjects. That is, it rests on the assumption that the data were generated by the same model for all participants, and that data are conditionally independent over subjects. This is known as fixed effects (FFX) inference, and it is not as robust to outliers as random effects (RFX) inference, which does not assume that the data were necessarily generated by the same model for each participant (Stephan et al., 2009).

Stephan et al. (2009) developed a novel VB approach for group level methods of Bayesian model comparison that used random effects instead of fixed effects analysis at the group level. They did this by treating models as random variables whose probabilities can be described by a Dirichlet distribution (which is conjugate to the multinomial distribution) with parameters that are estimated using the log-model evidences over all models and subjects (as described below). Once the optimal Dirichlet parameters have been estimated, they can be used to calculate posterior probabilities or exceedance probabilities of a given model for a randomly-selected participant. This is what is done in the third script (“BMS3_PPMs.m”, described in the Implementation section below), and the underlying mathematics is explained briefly below.

In the RFX approach introduced by Stephan et al. (2009), we assume that the probabilities of the different models (or hypotheses) are described by the following Dirichlet distribution:

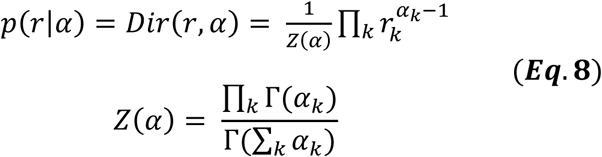

where *r* represents the probabilities *r* = [*r*_1_, …., *r*_K_] of *K* different models (or hypotheses), and *α* = [*α*_1_, …, *α_K_*] are related to unobserved “occurrences” of models in the population. This distribution is part of a hierarchical model: the next level depends on model probabilities, *r*, which are described by the Dirichlet distribution.

In the next level of the hierarchical model, we assume that the probability that a particular model generated the data of a particular subject, is given by a multinomial variable *m_n_* whose probability distribution is as follows:

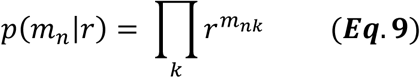

where *m_n_* is the multinomial variable that describes the probability that model *k* generated the data of subject *n* given the probabilities *r*.

Finally, in the lowest level of this hierarchical model, the probability of the data in the *n*th subject, given model *k*, over all parameters (ϑ) of the selected model (i.e. the marginal likelihood of the data in the nth subject, obtained by integrating over the parameters of the model) is given by:

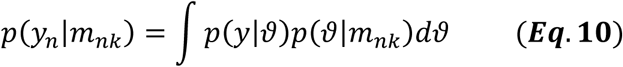

The goal is to invert this hierarchical model, that is, work backwards from data (*y*_n_) find the parameters of the Dirichlet distribution (which then allows the calculation of the expected posterior probability of obtaining the *k*th model for any randomly selected subject, as shown below). This model inversion is done using a VB approach in which the Dirichlet distribution is approximated with a conditional density, *q(r) = Dir(r, α*). Stephan et al. (2009) show that the following algorithm yields the optimal parameters of the conditional density *q(r) = Dir(r, α):*

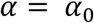

Until convergence

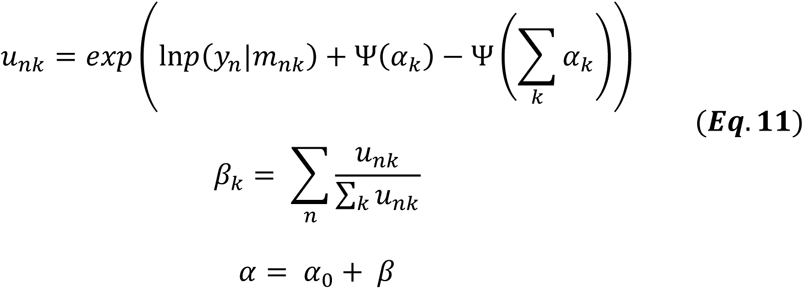

endwhere *α* are “occurrences” of models in the population; *α*_0_ is the Dirichlet prior, which, on the assumption that no models have been “seen” *a priori*, is set as *α*_0_ = [1,…,1] so that all models are equally probable to begin with; *u_nk_* is the non-normalised belief that model *k* generated the data *y_n_* for subject *n* (for the derivation of this line, please see Stephan et al., 2009); Ψ is the digamma function 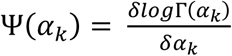; *β_k_* is the expected number of subjects whose data are believed to be generated by model *k* (so-called “data counts”); and the last line, *α = α*_0_ *+ β* essentially obtains the parameters of the Dirichlet distribution by starting with the Dirichlet prior *α*_0_ and adding on “data counts” *β* (Stephan et al., 2009).

Once the Dirichlet parameters have been optimised as per the algorithm above, this can be used for model comparisons at the group level. One way of comparing models is to simply compare the parameter estimates *α*. Another way is to calculate the multinomial parameters, 〈*r_k_*〉, that encode the posterior probability of model *k* being selected for a randomly chosen subject in the group:

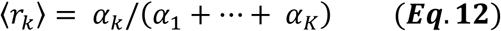

where *r_k_* is the probability of the model; the numerator of the fraction, *α_k_*, is the “occurrence” of model *k*; and the denominator (*α*_1_ *+* ⋯ + *α_K_*) is the sum of all model “occurrences.” This was how the PPMs were generated in the third script (“BMS3_PPMs.m”) below.

Another option for comparing models after the optimal Dirichlet parameters have been found, is to calculate the exceedance probability for a given model, as follows:

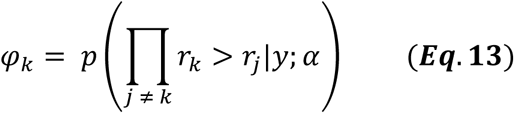

where φ*_k_* is the exceedance probability for model *k*, that is, the probability that it is more likely than any of the other models considered; *r_k_* is the probability of model *k*; *r_j_* is the probability of all other models considered; *y* represents the data and *α* represents the Dirichlet parameters.

Having introduced this RFX approach to VB, Stephan and colleagues (2009) then used both simulated and empirical data to demonstrate that when groups are heterogeneous, fixed effects analyses fail to remain sufficiently robust. Crucially, they also showed that RFX is robust to outliers, which can confound inference under FFX assumptions, when those assumptions are violated. Stephan et al. thus concluded that although RFX is more conservative than FFX, it is still the best method for selecting among competing neurocomputational models.

## 3 Methods

### 3.1 Experimental design

This experiment is a direct replication of that performed by Garrido et al. (2017), apart from the omission of a ‘divided attention’ condition. As they describe in greater detail in their paper, Garrido et al. (2017) utilised a novel audio-spatial attention task during which attention and prediction were orthogonally manipulated; this was done to evaluate the effect of surprise and attention in auditory processing (Garrido et al., 2017). The authors compared two models (shown in Figure 1) which may explain the effect attention has on the neural responses elicited by predicted and unpredicted events.

**Figure 1:**
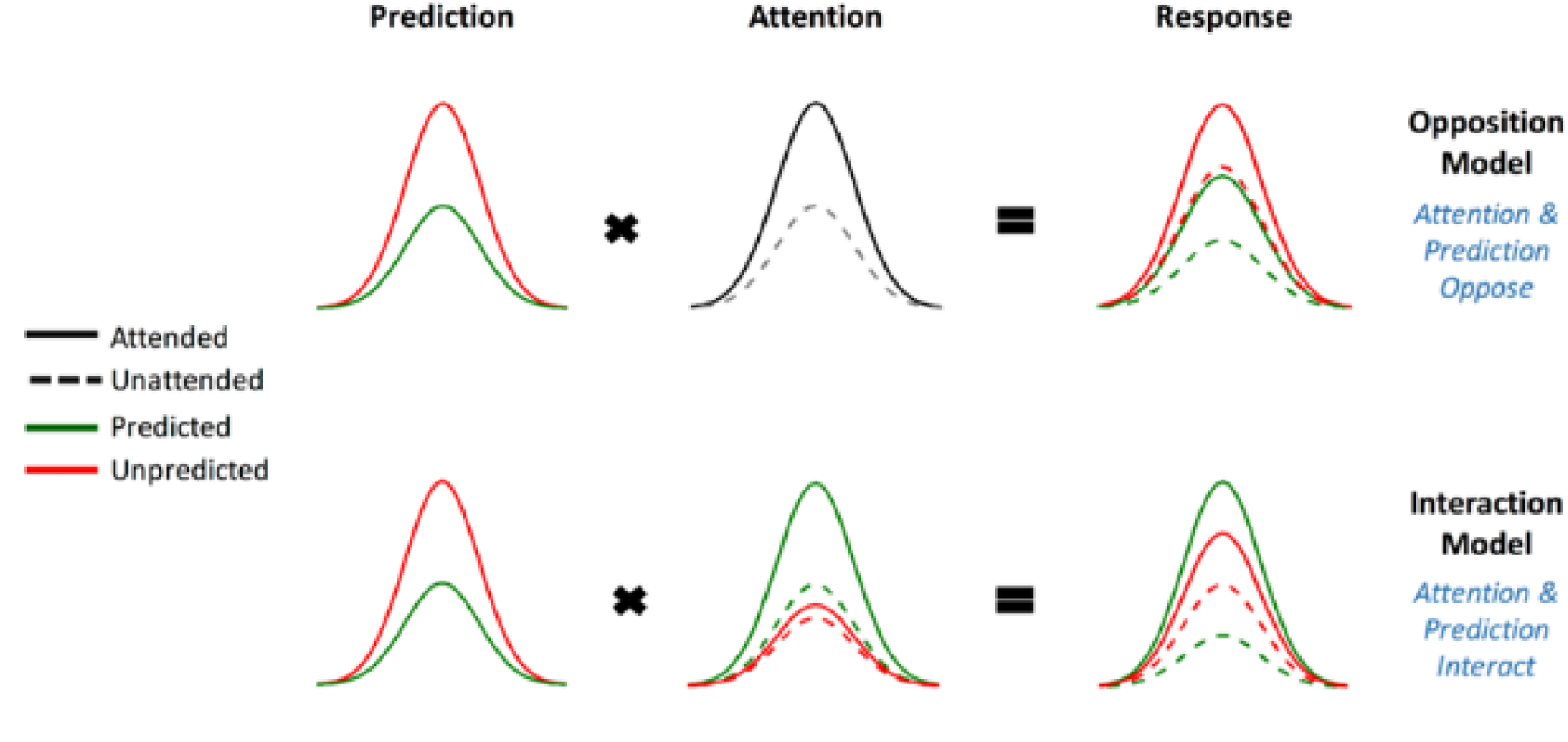
The two competing models that were evaluated using BMS. Reprinted with permission from Garrido et al. (2017) DOI: 10.1093/cercor/bhx087. Figure Published by Oxford University Press. All rights reserved. Available online at: https://academic.oup.com/cercor/advance-article/doi/10.1093/cercor/bhx087/3571164?searchresult=1. This figure is not covered by the Open-Access licence of this publication. For permissions contact

The original study supported the model in which attention boosts neural responses to both predicted and unpredicted stimuli, called the Opposition Model (Garrido et al., 2017). Prediction attenuates neural activity, while attention enhances this activity. Since these effects occur in opposite directions or have opposing effects, the researchers named the model (describing these effects) the Opposition Model. According to this model, attention improves the accuracy of predictions by precision weighting prediction errors more heavily. Thus, in light of this model, attention and prediction work together (in opposite directions) to improve our ability to make more accurate representations of the sensorium.

Our current study attempted to replicate the above-mentioned study with an independent dataset and employing the Bayesian methods that resembled the original study as closely as possible. The only difference was that the divided-attention condition was not administered because it was not required for the implementation and description of the BMS steps. It is hoped that the detailed description of our methods, adapted from those originally developed for fMRI by Rosa et al. (2010), prove to be useful for other EEG and/or MEG researchers. Furthermore, a replication study such as this one has the additional benefit of being responsive to the persisting replication crisis that continues to pose a significant problem for neuroscience and psychology (Hartshorne, 2012; Larson and Carbine, 2017; Szucs et al., 2017).

To this end we employed BMS to adjudicate between two competing hypotheses (see Figure 1), namely:

1. Attention increases (boosts) neural responses to both predicted and unpredicted stimuli. This is formalised in the Methods section and is then called Model One - the Opposition Model.
2. Attention boosts neural responses to predicted stimuli more than it boosts responses to unpredicted stimuli. This causes predicted attended stimuli to generate the highest neural responses, followed by attended unpredicted stimuli. This is formalised in the Methods section and is then called Model Two – the Interaction Model.

### 3.2 Participants

Twenty-one healthy adults (aged between 19-64 years, M = 25.00 years, SD= 9.83, nine females) were recruited via the University of Queensland’s Psychology Research Participation Scheme (SONA). Exclusion criteria included any history of mental or neurological disease, any previous head injury resulting in unconsciousness, or an age outside the prescribed range (18-65 years). All participants gave both written and verbal informed consent to both the study and to having their de-identified data made available in publicly distributed databases. Participants completed practice blocks of stimulus presentation prior to undergoing the EEG recording, in order to enable them to withdraw if they found the task unpleasant or excessively challenging. (No participants wished to withdraw.) Participants were monetarily compensated for their time. This study was approved by the University of Queensland Human Research Ethics Committee.

### 3.3 Task description

Participants wore earphones with inner-ear buds (Etymotic, ER3) and were asked to follow instructions on a computer screen. Participants were asked to pay attention to the sound stream in either the left or the right ear (ignoring the sounds that were being played in the other ear). Gaussian white noise was played to both ears and an oddball sequence was played to one of the ears. During a given block, participants were tasked with listening carefully for gaps in the white noise on the side to which they had been asked to attend. They were asked to press a “1” on the numbered keyboard when they heard a single gap (lasting 90 ms) in the white noise, and a “2” when they heard a double gap (two 90 ms gaps separated by 30 ms of white noise). They were asked to ignore any tones played on both the attended and the opposite ear. This task is described in further detail, including pictorial representations, in Garrido et al., (2017).

Participants listened to eight different blocks, each 190 seconds in duration. Each block contained a total of 30 targets (15 single gaps and 15 double gaps, randomly distributed across the block, but never occurring within 2.5 seconds of each other and never occurring at the same time as a tone). Throughout each block there were also 50-ms-long pure tones being played in one of the ears, with a 450 ms inter-stimulus interval. In each block there were two tones: the standard tone (either 500 Hz or 550 Hz counterbalanced between blocks) that occurred 85% of the time, and the deviant (either 550 Hz or 500 Hz, the opposite of the standard tone and counterbalanced across blocks) that occurred 15% of the time. All sound files were created using MATLAB (RRID:SCR_001622; The MathWorks, Inc.; http://www.mathworks.com) with sound recordings done using Audacity ^®^ (Audacity: Free Audio Editor and Recorder, RRID:SCR_007198) as previously described by Garrido et al., (2017). The order was counterbalanced such that no two participants received the same order of blocks.

Prior to and during the practice block/s, the volume of sound delivery was adjusted until the participant stated that they were able to hear the white noise well enough to complete the task. For each participant, an accuracy level was calculated, consisting of the percentage of white noise gaps that were correctly identified (as single or double) and responded to promptly (i.e. within two seconds of the gap/s). This was calculated separately for the practice block, which was repeated if a participant did not achieve at least 50% accuracy. Once participants achieved above 50% accuracy, they were invited to participate in the rest of the experiment. At the end of the experiment each participant’s accuracy was again calculated to ensure their accuracy level remained at least 50% (otherwise they were excluded from the study). This was to ensure that participants were attending to the task as instructed.

### 3.4 EEG data acquisition

Using a standardised nylon head cap fitted tightly and comfortably over the scalp, 64 silver/silver chloride (Ag/AgCl) scalp electrodes were placed according to the international 10-10 system for electrode placement. As is usual for this system, electrodes were placed above and below the left eye and just lateral to the outer canthi of both left and right eyes, to generate the vertical electrooculogram (VEOG) and horizontal electrooculogram (HEOG) recordings respectively. Continuous EEG data were recorded using a Biosemi Active Two system at a sampling rate of 1024 Hz. The onset of targets, standards and deviants were recorded with unique trigger codes at the time of delivery to the participant. Within each block, the target triggers were used for accuracy calculations, while the standard and deviant triggers were kept as the time points around which to epoch the data at a later stage.

### 3.5 EEG preprocessing

Following the collection of the raw EEG data, preprocessing was completed using Statistical Parametric Mapping (SPM) software (SPM12, RRID:SCR_007037; Wellcome Trust Centre for Neuroimaging, London; http://www.fil.ion.ucl.ac.uk/spm/). EEG data preprocessing included referencing data to the common average of all electrodes; downsampling to 200 Hz; bandpass filtering (between 0.5 to 40 Hz); eyeblink correction to remove trials marked with eyeblink artefacts (measured with the VEOG and HEOG channels); epoching using a peri-stimulus window of −100 to 400 ms; artefact rejection (with 100 uV cut-off); low-pass filtering (40 Hz; to remove any high frequency noise from the robust averaging step) and baseline correction (−100 to 0 ms window).

#### Source Reconstruction

For source BMS, SPM12 software was used to obtain source estimates on the cortex by reconstructing the scalp activity using a single-shell head model. The forward model was then inverted with multiple sparse priors (MSP) assumptions for the variance components (Friston et al., 2008) under group constraints (Litvak and Friston, 2008). The entire time window of 0 to 400 ms was used to infer the most likely cortical regions that generated the data observed during this time. Images for each participant and each condition were obtained from the source reconstructions and were smoothed at full width at half maximum (FWHM) 12 x 12 x 12 mm. This source reconstruction step is available as an online script (named “BMS1_Source_ImCreate.m” and available at https://github.com/ClareDiane/BMS4EEG).

### 3.6 Bayesian Model Selection Maps: Implementation for M/EEG

For all data analysis steps (Table 1), we used SPM12 software package for MATLAB. We wrote inhouse MATLAB scripts, integrated with SPM12 and now available online (https://github.com/ClareDiane/BMS4EEG). The online scripts are divided into three BMS scripts. In the first script (BMS1_ST_ImCreate.m for spatiotemporal BMS and BMS1_Source_ImCreate.m for source BMS), we call the preprocessed EEG data and then create images for every trial, for every condition, and for every participant. The second script (BMS2_ModelSpec_VB.m) specifies the hypotheses and implements variational Bayes (as described in the Theory section). The last script (BMS3_PPMs.m) then creates Posterior Probability Maps.

In the model specification and VB script (BMS2_ModelSpec_VB.m), we changed individual participants’ data file structures in order to match the format that SPM typically requires to read fMRI data. This is done by first loading the relevant file path and then changing the file structure. Once these newly-structured files had been saved, we next specified the models to be compared: this was done by assigning covariate weights to describe both models (please see the instructions contained within BMS2_ModelSpec_VB.m on Github). The Opposition Model was assigned weights of [1, 2, 2, and 3] for the unattended predicted, attended predicted and unattended unpredicted, and attended unpredicted, respectively. The Interaction Model was assigned weights of [1, 4, 2, and 3] for the same conditions.

These covariate weights essentially describe the assumed relationship between the different conditions according to a given model. For example, using [1, 2, 2, and 3] as employed in the Opposition Model, means that according to the Opposition Model, the unattended predicted condition (the first condition with an assigned weight of 1) evokes the smallest activity, whereas the attended unpredicted (the fourth condition with a weight of 3) has the greater activity, and both attended predicted and unattended unpredicted (second and third conditions with an equal weight of 2) are in between the former two conditions and indistinguishable in magnitude from each other.

We then created log-evidence images, representing the log-model evidences, for both models (separately), for every participant (individually) at every voxel. In the case of spatiotemporal (scalp-level) BMS, each voxel was representative of a specific spatiotemporal location within the peristimulus time window (0 to 400 ms) and topologically across the scalp, such that the first two dimensions of the voxel refer to the space across the scalp and the third dimension is time (as shown in Figure 2). Conversely, in the source BMS (which began with the source reconstruction steps described above), each voxel was representative of an inferred location in three-dimensional source space. Once log evidence images had been created, these were smoothed with a 1 mm half-width Gaussian kernel.

In summary, one can create posterior probability maps or log evidence maps in sensor or source space. In sensor space, this involves creating a two-dimensional image over the scalp surface and equipping the space with a peristimulus time dimension. This creates posterior probability maps over the scalp surface and peristimulus time, enabling one to identify regionally and temporally specific effects due to a particular model, relative to other contrasts. Alternatively, one can create three-dimensional posterior probability maps in source space, following source reconstruction.

The core SPM script that allows VB to be used on fMRI data is named spm_spm_vb.m and is found in the SPM12 package, downloadable from the SPM site. This core script was edited in order to adapt the VB method for EEG, as follows. Changes were made such that different data structures could be read in the same way that fMRI data would usually be read. Furthermore, high-pass filtering steps were removed as these only apply to low-frequency drifts associated with fMRI data. The specific changes made between the original script and the altered one to be used for spatiotemporal BMS are accessible online (goo.gl/ZVhPT7). For the source BMS steps, the same changes were left in place as outlined above, and in addition, the required minimum cluster size was changed from 16 voxels to 0 voxels to allow for visualisation of all clusters of any size. The specific differences between the original and source BMS versions of the spm_spm_vb script are accessible online (goo.gl/WXAo67).

In the final step (BMS3_PPMs.m), the SPM Batch Editor was used to apply a random effects approach to the group model evidence data in a voxel-wise manner, thus translating the log-evidence images from the previous step into Posterior Probability Maps (similar to how Penny at al. (2007) and Rosa et al., (2010) have produced PPMs previously for fMRI data). The maps, displayed in the Figures 2, 3 and 4, were generated by selecting threshold probabilities of 75% for the spatiotemporal maps (Figure 2) and 50% for the source maps (Figures 3 and 4). This threshold can be adjusted by the user. EPMs can also be displayed by selecting the relevant setting in the final script (please see the instructions on Github).

**Table 1:** Step-by-step summary of method

| **Table 1: Step-by-step summary of method** |  |
| --- | --- |
| Task: | Suggested steps: |
| Saving the correct spm_spm_vb.m files | 1. Find and open the spm12 folder on your computer.<br>2. Find the spm_spm_vb.m script in that folder, and rename this to spm_spm_vb_fMRI.m. Then add the spm_spm_vb_ST.m and spm_spm_vb_source.m scripts (saved in the associated Github repository) to your spm12 folder.<br>3. Before undertaking either the spatiotemporal BMS or source BMS steps, rename the currently-relevant script from the above step to spm_spm_vb.m. Once you have finished the BMS steps, rename the script back to its original name, to re-identify it as being for either the spatiotemporal ('spm_spm_vb_ST.m') or source BMS ('spm_spm_vb_source.m'). In this way, you will keep track of which spm_spm_vb.m script to use for whichever BMS steps you are about to do. |
| Creating spatiotemporal ("scalp") PPMs: | 1. BMS script 1: Change the file paths to reflect the location of ERP data.<br>2. Run BMS script 1: BMS1_ST_ImCreate.m.<br>3. Ensure the correct spm_spm_vb.m file is saved in SPM12 folder.<br>4. Run BMS script 2: BMS2_ModelSpec_VB.m.<br>5. Run BMS script 3: BMS3_PPMs.m. Threshold is set to 0.75 and adjustable. |
| Creating source PPMs: | 1. BMS script 1: Change the file paths to reflect location of source reconstructed images.<br>2. Run BMS script 1: BMS1_Source_ImCreate.m.<br>3. Ensure the correct spm_spm_vb.m file is saved in SPM12 folder.<br>4. Run BMS script 2: BMS2_ModelSpec_VB.m.<br>5. Replace NaNs with zeros in the LogEv.nii files: BMS2b_Source_NaNtoZeros.m.<br>6. Run BMS script 3: BMS3_PPMs.m. Adjust probability threshold as desired. |

## 4 Results

The raw dataset for this study can be found on Figshare (EEG_Auditory_Oddball_Raw_Data repository, https://figshare.com/s/1ef6dd4bbdd4059e3891).

The preprocessed dataset for this study can also be found on Figshare (EEG_Auditory_Oddball_Preprocessed_Data repository, https://figshare.com/s/c6e1f9120763c43e6031).

### 4.1 Scalp - Spatiotemporal

Figure 2 shows scalp (spatiotemporal) PPMs of the two competing models over space and time. These maps display all posterior probabilities exceeding 75% over space and time for both models. As can be seen in the figure, spatiotemporal BMS results revealed that Model One (the Opposition Model) was by and large the superior model. The Opposition Model had model probabilities exceeding 75% across the majority of later time points (with most significant clusters between 225-360 ms), and over most frontocentral and bilateral channel locations, as shown in (**A**). On the other hand, as shown in (**B**), the Interaction Model did have over 75% model probability centrally between 175-185 ms, which is within the mismatch negativity (MMN) time window. These findings replicate those of Garrido et al., (2017), and strongly support the implications discussed in great depth in that paper.

**Figure 2:**
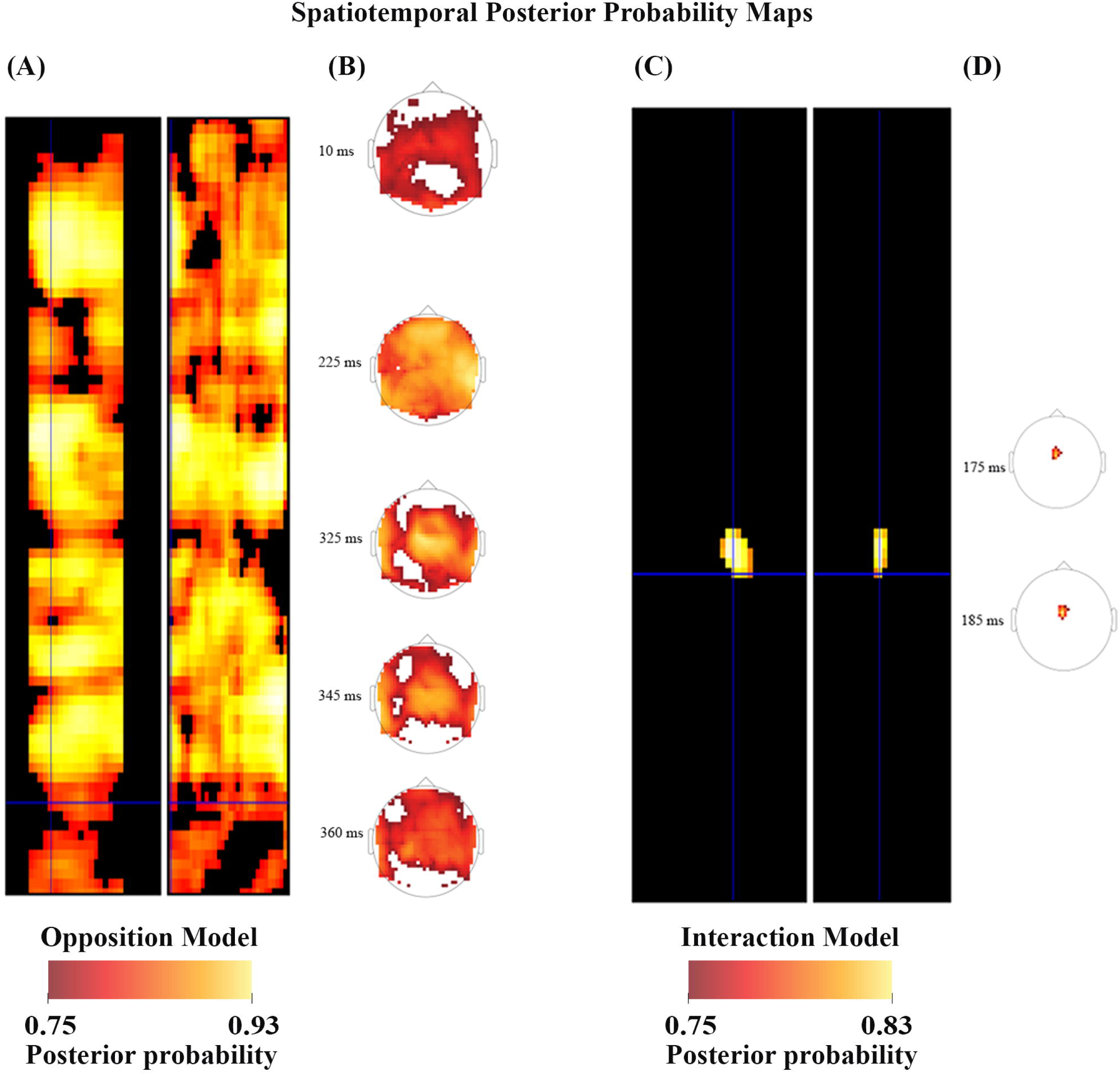
Scalp Posterior Probability Maps of the two competing models over space and time. (The scalp images include the participant’s nose, pointing upwards, and ears, visible as if viewed from above.) These maps display all posterior probabilities exceeding 75% over space and time for both models. The left sides of both panels (**A**) and (**C**) both depict the temporal information, showing the model probabilities at each point in time from 0 ms (when the tone was played, at the top of the diagrams) to 400 ms after the stimulus presentation (at the bottom of the diagram), across the surface of the scalp (which traverses the width of the panels). The right sides (**B**) and (**D**) show the spatial locations of the probability clusters which exceeded the threshold of 75% probability.

### 4.2 Source

As shown in Figures 3, 4 and 5, source BMS results also favoured the Opposition Model, with higher model probability over the left supramarginal gyrus (with 91% model probability over a relatively large cluster, K_E_ = 6115), the right superior temporal gyrus (with 87% model probability over a cluster with K_E_ = 5749) as well as over parts of the left inferior parietal lobe, right inferior parietal lobe and left postcentral gyrus. Having said this, the Interaction Model also had two large clusters, albeit with lower model probabilities compared to the Opposition Model’s highest-probability clusters: specifically, the Interaction Model had a cluster of size K_E_ = 6346 over the left inferior parietal lobe and a cluster of size K_E_ = 5353 over the right inferior parietal lobe (with 74% model probability in both places).

**Figure 3:**
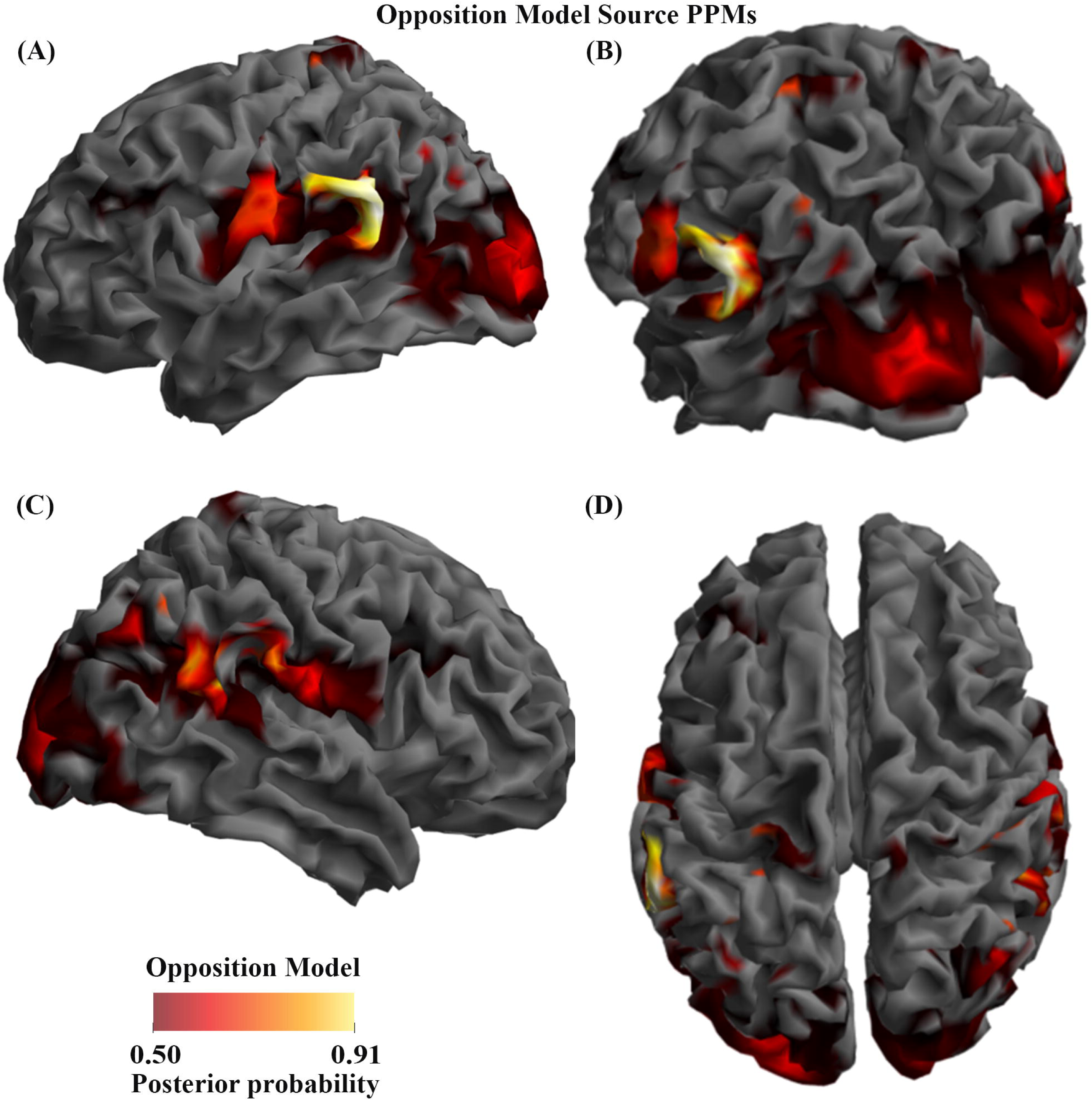
Source Posterior Probability Map for the Opposition Model (that is, reconstructed images representing the model inference at the group level for this model), thresholded at >50% posterior probability. (**A**): view from the left side. (**B**): view from the left side, from the posterior (back) end. (**C**): view from the right side. (**D**): view from above.

**Figure 4:**
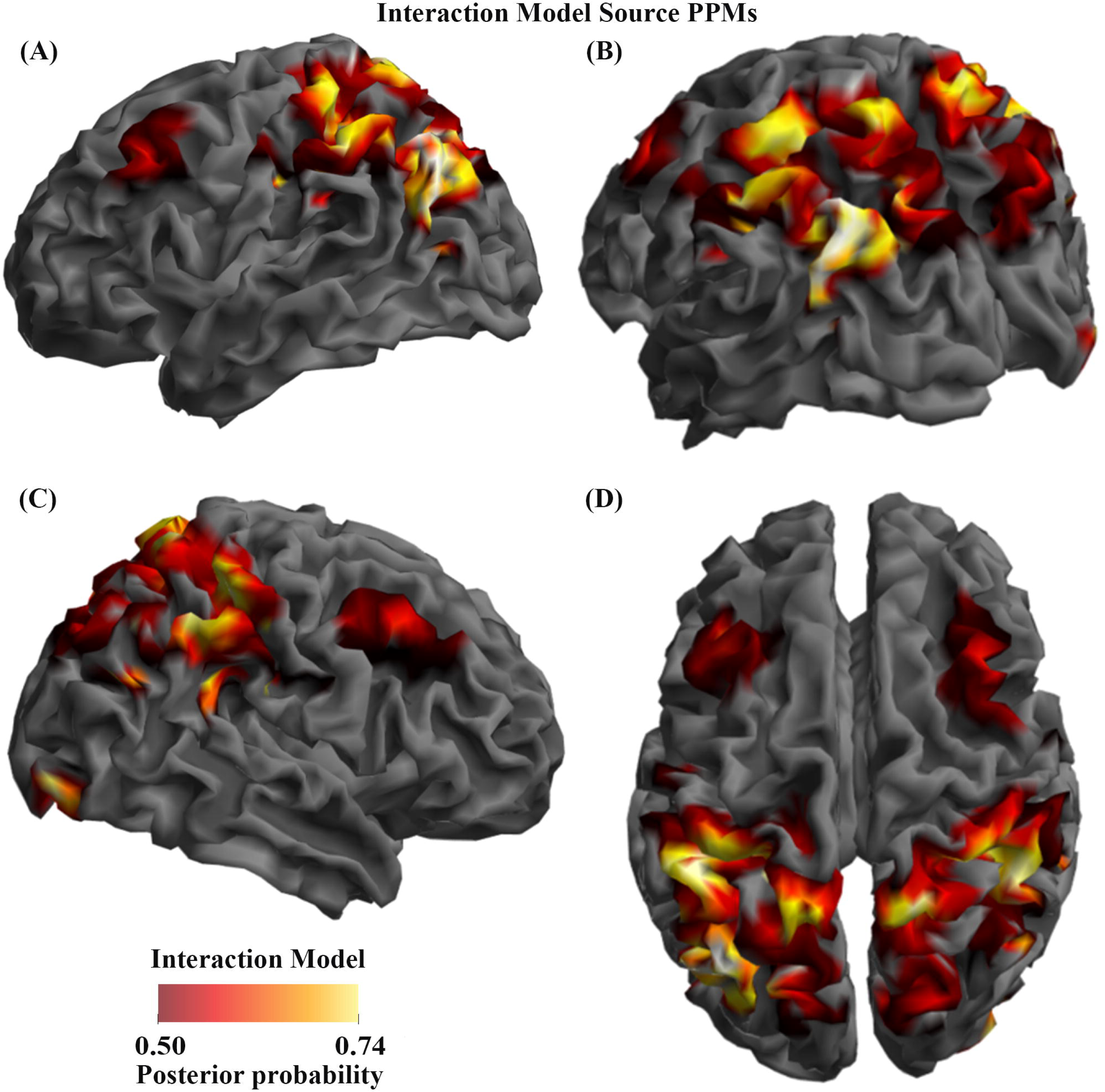
Source Posterior Probability Map for the Interaction Model (that is, reconstructed images representing the model inference at the group level for this model), thresholded at >50% posterior probability. (**A**): view from the left side. (**B**): view from the left side, from the posterior (back) end. (**C**): view from the right side. (**D**): view from above.

**Figure 5:**
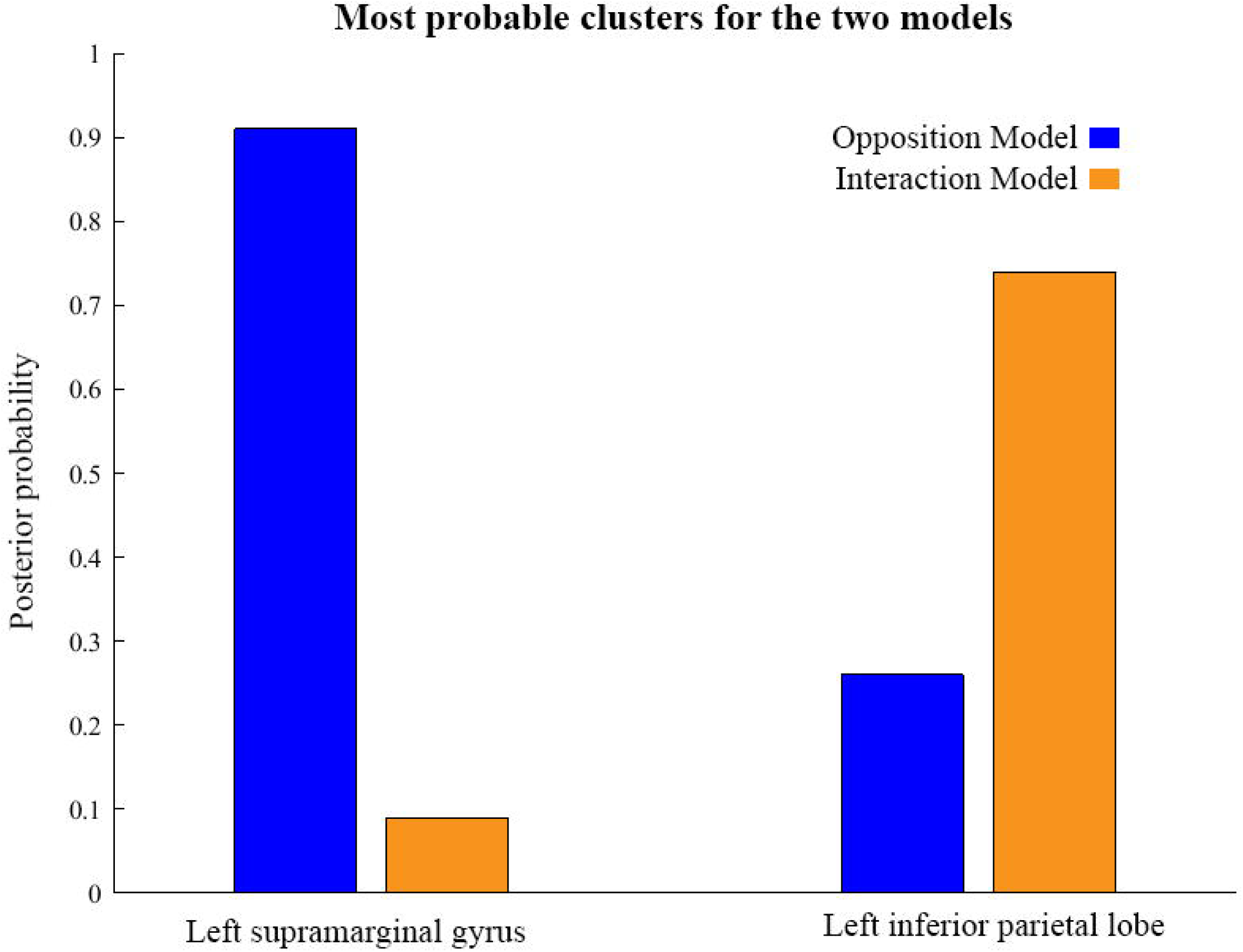
Comparison of the posterior probabilities for the two models at the location of the highest-probability cluster of the Opposition Model (left) and the location of the highest-probability cluster of the Interaction Model (right). The left supramarginal gyrus cluster, which was the highest probability cluster for the Opposition Model (left), was located at Montreal Neurological Institute (MNI) coordinates (62, −42, 30), while the left inferior parietal lobe cluster, which was the highest probability cluster for the Interaction Model, was located at MNI coordinates (−54, −32, 46).

Figures 3 and 4 show that different brain regions are likely to perform different computations best described by the Opposition and Interaction Models, respectively. Furthermore, Figure 5 compares the magnitude of the calculated posterior probabilities, at the locations of the highest probability cluster for both models. The possible functional reasons for the different anatomical locations that emerge for the two different models may be an interesting subject for future study, but fall outside the scope of this methods paper. In any case, the usefulness of this probability mapping approach illustrated in Figures 2, 3 and 4, lies in the ability of pinpointing where and when given computations are likely to be performed in the brain.

## 5 Discussion

This paper shows how to use RFX Bayesian model selection mapping methods for M/EEG data analysis. This method was originally developed for fMRI by Rosa and colleagues (2010), and provides a way of displaying the probabilities of different cognitive models at different timepoints and brain locations, given a neuroimaging dataset. We aimed to provide an in-depth explanation, written in a didactical manner, of the BMS and posterior probability mapping steps that were successfully used by Garrido et al. (2017) in their recent EEG paper.

Being a Bayesian approach to hypothesis-testing, the method described here provides multiple advantages over frequentist inference methods. The first of these advantages is that VB allows for comparisons between non-nested models. Consequently, it is especially useful in the context of model-based neuroimaging (Montague et al., 2004; O’Doherty et al., 2007; Rosa et al., 2010; Garrido et al., 2017). Another advantage is that the probability of the null hypothesis itself can be assessed (instead of simply being, or failing to be, rejected). A final advantage is that, although only two models were compared here, the same method can also be applied to any arbitrary number of models. For example, the analyses described here could proceed slightly differently, based on the same data but introducing another (or multiple other) model/s against which to compare the Opposition and Interaction Models. Potentially, any number of theoretically motivated models could be considered. Considering all of these advantages, the method described here should prove useful in a wide variety of M/EEG experiments.

In summary, we have shown here how to adapt Bayesian Model Selection maps, originally developed for fMRI data by Rosa and colleagues (2010), to M/EEG data analysis. It is hoped that the reporting of analytical methods such as these, as well as the availability of all the code and dataset, will not only contribute to the Open Science movement, but may also encourage other researchers to adopt this novel M/EEG data analysis method in a way that is useful for addressing their own neuroscience questions. We postulate that the use of this Bayesian model mapping of M/EEG data to adjudicate between competing computational models in the brain, both at the scalp and source level, will be a significant advancement in the field of M/EEG neuroimaging and may provide new insights in cognitive neuroscience.

## 7 Author Contributions

MG designed the study and the analysis methods. ER wrote the code. CH and RR collected and analysed the data, and organised the data and code for sharing. CH wrote the first draft of the manuscript. ER, RR and MG edited the manuscript.

## 9 Acknowledgments

The authors thank the participants for their time. The authors are grateful to Maria Rosa for helpful advice regarding source BMS steps, and to the two anonymous reviewers, whose helpful comments and suggestions led to significant improvements in the paper. The authors also thank Jessica McFadyen for help with preprocessing, Ilvana Dzafic for help with EEG data acquisition, and Jeremy Taylor for sharing tools for visualising spatiotemporal images (shown in panels (**B**) and (**D**) in Figure 2).

## 11 Supplementary Material

Please see the Supplementary Material attached.

## References

Adams, R. A., Friston, K. J., & Bastos, A. M. (2015). Active inference, predictive coding and cortical architecture. In Recent Advances on the Modular Organization of the Cortex (pp. 97–121). Springer Netherlands.

Akaike, H. (1980). Likelihood and the Bayes procedure. Trabajos de estadística y de investigación operativa, 31(1), 143–166.

Anllo-Vento, L. (1995). Shifting attention in visual space: the effects of peripheral cueing on brain cortical potentials. The International journal of neuroscience 80(1–4): 353–370.

Arnott, R. S., and R. C. Alain (2002). Stepping out of the spotlight: MMN attenuation as a function of distance from the attended location. NeuroReport 13(17): 2209–2212.

Auksztulewicz, R. and K. Friston (2015). Attentional Enhancement of Auditory Mismatch Responses: a DCM/MEG Study. Cerebral Cortex 25(11): 4273–4283.

Bayes, T & Price, R. (1763). An essay towards solving a problem in the doctrine of chances. by the late Rev. Mr. Bayes, frs communicated by Mr. Price, in a letter to John Canton, amfrs. Philosophical Transactions (1683–1775), 370–418.

Beal, M., Ghahramani, Z., 2003. The variational Bayesian EM algorithms for incomplete data: with application to scoring graphical model structures. In: Bernardo, J., Bayarri, M., Berger, J., Dawid, A. (Eds.), Bayesian Statistics 7. Cambridge University Press.

Boly, M., M. Garrido, O. Gosseries, M.-A. Bruno, P. Boveroux, C. Schnakers, M. Massimini, V. Litvak, S. Laureys and K. Friston (2011). Preserved Feedforward But Impaired Top-Down Processes in the Vegetative State. Science (Washington) 332(6031): 858–862.

Carter, T., & Fe, S. (2007). An introduction to information theory and entropy. Complex systems summer school, Santa Fe.

Chaumon, M., V. Drouet and C. Tallon-Baudry (2008). Unconscious associative memory affects visual processing before 100 ms. Journal of Vision 8(3): 10.1–10.

Chun, M. M. and Y. Jiang (1998). Contextual Cueing: Implicit Learning and Memory of Visual Context Guides Spatial Attention. Cognitive Psychology 36(1): 28–71.

Connor, J. T., Elm, J. J., Broglio, K. R., & ESETT and ADAPT-IT Investigators. (2013). Bayesian adaptive trials offer advantages in comparative effectiveness trials: an example in status epilepticus. Journal of clinical epidemiology, 66(8), S130–S137.

Dayan, P., Hinton, G. E., Neal, R. M., & Zemel, R. S. (1995). The helmholtz machine. Neural computation, 7(5), 889–904.

Dienes, Z. (2016). How Bayes factors change scientific practice. Journal of Mathematical Psychology 72: 78–89.

Doherty, J., A. Rao, M. Mesulam and A. Nobre (2005). Synergistic Effect of Combined Temporal and Spatial Expectations on Visual Attention. Journal of Neuroscience 25(36): 8259–8266.

Dunson, D. B. (2001). Commentary: practical advantages of Bayesian analysis of epidemiologic data. American journal of Epidemiology, 153(12), 1222–1226.

Feldman, H., & Friston, K. (2010). Attention, uncertainty, and free-energy. Frontiers in human neuroscience, 4, 215–238.

Feynman, R. P., & Brown, L. M. (1942). Feynman’s thesis: a new approach to quantum theory. World Scientific.

Feynman, R. P., Hibbs, A. R., & Styer, D. F. (2010). Quantum mechanics and path integrals. Courier Corporation.

Friston K, Penny W (2003) Posterior probability maps and SPMs. NeuroImage 19: 1240–1249.

Friston, K. (2007a). APPENDIX 1 - Linear models and inference. Statistical Parametric Mapping. London, Academic Press: 589–591.

Friston, K. (2007b). CHAPTER 1 - A short history of SPM. Statistical Parametric Mapping. London, Academic Press: 3–9.

Friston, K. (2012). The history of the future of the Bayesian brain. NeuroImage 62(2): 1230–1233.

Friston, K., Harrison, L., Daunizeau, J., Kiebel, S., Phillips, C., Trujillo-Barreto, N., Henson, R., Flandin, G., Mattout, J., 2008. Multiple sparse priors for the M/EEG inverse problem. NeuroImage, 39(3), 1104–1120.

Friston, K. and W. Penny (2007). CHAPTER 22 - Empirical Bayes and hierarchical models. Statistical Parametric Mapping. London, Academic Press: 275–294.

Friston, K. and W. Penny (2007). CHAPTER 23 - Posterior probability maps. Statistical Parametric Mapping. London, Academic Press: 295–302.

Friston, K. J., N. Trujillo-Barreto and J. Daunizeau (2008). DEM: A variational treatment of dynamic systems. Neuroimage 41(3): 849–885.

Friston, K., S. Kiebel, M. Garrido and O. David (2007). CHAPTER 42 - Dynamic causal models for EEG. Statistical Parametric Mapping. London, Academic Press: 561–576.

Garrido, M. I., C. L. J. Teng, J. A. Taylor, E. G. Rowe and J. B. Mattingley (2016). Surprise responses in the human brain demonstrate statistical learning under high concurrent cognitive demand. npj Science of Learning 16006: 1–7.

Garrido, M. I., M. Sahani and R. J. Dolan (2013). Outlier Responses Reflect Sensitivity to Statistical Structure in the Human Brain (Statistical Learning and Outlier Detection). 9(3): e1002999: 1–10.

Garrido, M., Rowe, E., Halasz, V., & Mattingley, J. (2017). Bayesian mapping reveals that attention boosts neural responses to predicted and unpredicted stimuli. Cerebral Cortex, 1–12.

Hartshorne, J. (2012). Tracking replicability as a method of post-publication open evaluation. Frontiers in Computational Neuroscience 6(NA): 70–83.

Hohwy, J. (2013). The predictive mind. Oxford University Press.

Kane, N. M., S. H. Curry, S. R. Butler and B. H. Cummins (1993). Electrophysiological indicator of awakening from coma. 341: 688–688.

Kass, R. E., & Raftery, A. E. (1995). Bayes factors. Journal of the american statistical association, 90(430), 773–795.

Kok, P., D. Rahnev, J. F. M. Jehee, H. C. Lau and F. P. de Lange (2012). Attention Reverses the Effect of Prediction in Silencing Sensory Signals. Cerebral Cortex 22(9): 2197–2206.

Kujala, T. and R. Näätänen (2010). The adaptive brain: A neurophysiological perspective. Progress in Neurobiology 91(1): 55–67.

Kullback, S., & Leibler, R. A. (1951). On information and sufficiency. The annals of mathematical statistics, 22(1), 79–86.

Lappalainen, H., Miskin, J.W., 2000. Ensemble Learning, in: Girolami, M. (Ed.), Advances in Independent Component Analysis. Springer-Verlag, Berlin.

Larson, M. J. and K. A. Carbine (2017). Sample size calculations in human electrophysiology (EEG and ERP) studies: A systematic review and recommendations for increased rigor. International Journal of Psychophysiology 111: 33–41.

Lieder, F., J. Daunizeau, M. Garrido, K. Friston and K. Stephan (2013). Modelling Trial-by-Trial Changes in the Mismatch Negativity. PLoS Computational Biology 9(2) e1002911: 1–16.

Litvak, V., Friston, K., 2008. Electromagnetic source reconstruction for group studies, NeuroImage, pp. 1490–1498.

Lv, J.-Y., T. Wang, J. Qiu, S.-H. Feng, S. Tu and D.-T. Wei (2010). The electrophysiological effect of working memory load on involuntary attention in an auditory–visual distraction paradigm: an ERP study. Experimental Brain Research 205(1): 81–86.

Mangun, G. R. and S. A. Hillyard (1991). Modulations of Sensory-Evoked Brain Potentials Indicate Changes in Perceptual Processing During Visual–Spatial Priming. Journal of Experimental Psychology: Human Perception and Performance 17(4): 1057–1074.

Meinert, C. L. (2012). Frequentist vs. Bayesian Analysis. Hoboken, NJ, USA, Hoboken, NJ, USA: John Wiley & Sons, Inc.

Mohammad-Djafari, A. (2002). Bayesian inference for inverse problems. AIP Conference Proceedings, 617(1), 477–496.

Montague, P.R., Hyman, S.E., Cohen, J.D., Oct 2004. Computational roles for dopamine in behavioural control. Nature 431, 760–767.

Näätänen, R. and K. Alho (1997). Higher-order processes in auditory-change detection. Trends in Cognitive Sciences 1(2): 44–45.

Näätänen, R., T. Kujala and I. Winkler (2011). Auditory processing that leads to conscious perception: A unique window to central auditory processing opened by the mismatch negativity and related responses. Psychophysiology 48(1): 4–22.

Naeaetaenen, R., M. Tervaniemi, E. Sussman, P. Paavilainen and I. Winkler (2001). 'Primitive intelligence’ in the auditory cortex. R. Naeaetaenen. 24: 283–288.

Neal, R. (1998). Annealed importance sampling (Technical Report 9805 (revised)). Department of Statistics, University of Toronto.

Needham, C. J., Bradford, J. R., Bulpitt, A. J., & Westhead, D. R. (2007). A primer on learning in Bayesian networks for computational biology. PLoS computational biology, 3(8), e129.

Neyman, J. and E. S. Pearson (1933). On the Problem of the Most Efficient Tests of Statistical Hypotheses. Philosophical Transactions of the Royal Society of London. Series A, Containing Papers of a Mathematical or Physical Character (1896–1934) 231(694): 289–337.

O’Doherty, J.P., Hampton, A., Kim, H., May 2007. Model-based fMRI and its application to reward learning and decision making. Ann. N.Y. Acad. Sci. 1104, 35–53.

Penny, W. D. (2012). Comparing dynamic causal models using AIC, BIC and free energy. NeuroImage, 59(1), 319–330.

Penny, W. D., & Ridgway, G. R. (2013). Efficient posterior probability mapping using Savage-Dickey ratios. PLoS One, 8(3), e59655.

Penny, W. D., J. Mattout and N. Trujillo-Barreto (2007). CHAPTER 35 - Bayesian model selection and averaging. Statistical Parametric Mapping. London, Academic Press: 454–467.

Penny, W. D., Trujillo-Bareto, N., & Flandin, G. (2005). Bayesian analysis of single-subject fMRI data: SPM5 implementation. Wellcome Trust Centre for Neuroimaging, UCL, London.

Penny, W. D., Trujillo-Barreto, N. J., & Friston, K. J. (2005). Bayesian fMRI time series analysis with spatial priors. NeuroImage, 24(2), 350–362.

Penny, W., & Flandin, G. (2005). Bayesian analysis of fMRI data with spatial priors. In Proceedings of the Joint Statistical Meeting (JSM). American Statistical Association.

Penny, W., & Sengupta, B. (2016). Annealed importance sampling for neural mass models. PLoS computational biology, 12(3), e1004797.

Penny, W., Flandin, G., & Trujillo-Barreto, N. (2007). Bayesian comparison of spatially regularised general linear models. Human brain mapping, 28(4), 275–293.

Penny, W., Kiebel, S., & Friston, K. (2003). variational Bayesian inference for fMRI time series. NeuroImage, 19(3), 727–741.

Penny, W., S. Kiebel and K. Friston (2007). CHAPTER 24 - variational Bayes. Statistical Parametric Mapping. London, Academic Press: 303–312.

Penny, W.D., Stephan, K.E., Mechelli, A., Friston, K.J., (2004). Comparing dynamic causal models. NeuroImage 22 (3), 1157–1172.

Rao, R. P. N. and D. H. Ballard (1999). Predictive coding in the visual cortex: a functional interpretation of some extra-classical receptive-field effects. Nature Neuroscience 2(1): 79–87.

Rosa, M., S. Bestmann, L. Harrison and W. Penny (2010). Bayesian Model Selection Maps for Group Studies. NeuroImage, 49(1), 217–224.

Rowe, E.G., Harris C.D., Randeniya, R. and Garrido, M.I. (2018). Bayesian Model Selection Maps for group studies using M/EEG data: EEG_Auditory_Oddball_Preprocessed_Data. Figshare. (2018) https://figshare.com/s/c6e1f9120763c43e6031 doi: 10.6084/m9.figshare.5812764

Rowe, E.G., Harris C.D., Randeniya, R. and Garrido, M.I. (2018). Bayesian Model Selection Maps for group studies using M/EEG data: EEG_Auditory_Oddball_Raw_Data. Figshare. (2018), https://figshare.com/s/1ef6dd4bbdd4059e3891 doi: 10.6084/m9.figshare.5769141

Sallinen, M., J. Kaartinen and H. Lyytinen (1994). Is the appearance of mismatch negativity during stage 2 sleep related to the elicitation of K-complex? Electroencephalography and Clinical Neurophysiology 91(2): 140–148.

Shannon, C. E. (2001 {originally 1948}). A mathematical theory of communication. ACM SIGMOBILE Mobile Computing and Communications Review, 5(1), 3–55.

Stephan, K. E., W. D. Penny, J. Daunizeau, R. J. Moran and K. J. Friston (2009). Bayesian model selection for group studies. NeuroImage 46(4): 1004–1017.

Summerfield, C. and T. Egner (2009). Expectation (and attention) in visual cognition. Trends in Cognitive Sciences 13(9): 403–409.

Summerfield, J. J., J. Lepsien, D. R. Gitelman, M. M. Mesulam and A. C. Nobre (2006). Orienting Attention Based on Long-Term Memory Experience. Neuron 49(6): 905–916.

Szucs, D., J. P. A. Ioannidis and E.-J. Wagenmakers (2017). Empirical assessment of published effect sizes and power in the recent cognitive neuroscience and psychology literature. PLoS Biology 15(3): 1–18.

Trippa, L., Lee, E. Q., Wen, P. Y., Batchelor, T. T., Cloughesy, T., Parmigiani, G., & Alexander, B. M. (2012). Bayesian adaptive randomized trial design for patients with recurrent glioblastoma. Journal of Clinical Oncology, 30(26), 3258–3263.

Vallverdú, J. (2008). The false dilemma: Bayesian vs. frequentist. arXiv preprint arXiv:0804.0486.

Woolrich M, Jenkinson M, Brady M, Smith S (2004) Fully Bayesian spatio-temporal modeling of fMRI data. IEEE Trans Med Imaging 23: 213–231.

Woolrich, M.W., Behrens, T.E., Beckmann, C.F., Smith, S.M., Jan 2005. Mixture models with adaptive spatial regularization for segmentation with an application to fMRI data. IEEE Trans. Med. Imaging 24, 1–11.

Yucel, G., C. Petty, G. McCarthy and A. Belger (2005). Graded Visual Attention Modulates Brain Responses Evoked by Task-irrelevant Auditory Pitch Changes. Journal of Cognitive Neuroscience 17(12): 1819–1828.

